# Proguanil and chlorhexidine augment the antibacterial activities of clarithromycin and rifampicin against *Acinetobacter baumannii*

**DOI:** 10.1101/2023.07.14.549121

**Authors:** Chuandong Wang, Tingting Zhang, Yan Wang, Yipeng Wang, Hongwei Pan, Xinyu Dong, Siyu Liu, Meng Cao, Shuhua Wang, Mingyu Wang, Yuezhong Li, Jian Zhang, Wei Hu

## Abstract

The emergence of *Acinetobacter baumannii* infections as a significant healthcare concern in hospital settings, coupled with their association with poorer clinical outcomes, has prompted extensive investigation into novel therapeutic agents and innovative treatment strategies. Proguanil and chlorhexidine, both categorized as biguanide compounds, have displayed clinical efficacy as antimalarial and topical antibacterial agents, respectively. In this study, we conducted an investigation to assess the effectiveness of combining proguanil and chlorhexidine with clarithromycin or rifampicin against both laboratory strains and clinical isolates of *A. baumannii*. The combination therapy demonstrated rapid bactericidal activity against planktonic multidrug-resistant *A. baumannii*, exhibiting efficacy in eradicating mature biofilms and impeding the development of antibiotic resistance *in vitro*. Additionally, when administered in conjunction with clarithromycin or rifampicin, proguanil enhanced the survival rate of mice afflicted with intraperitoneal *A. baumannii* infections, and chlorhexidine expedited wound healing in mice with skin infections. These findings are likely attributable to the disruption of *A. baumannii* cell membrane integrity by proguanil and chlorhexidine, resulting in heightened membrane permeability and enhanced intracellular accumulation of clarithromycin and rifampicin. Overall, this study underscores the potential of employing proguanil and chlorhexidine in combination with specific antibiotics to effectively combat *A. baumannii* infections and improve treatment outcomes in clinically challenging scenarios.

**IMPORTANCE:** *A. baumannii* has emerged as a globally significant nosocomial pathogen due to its remarkable ability to acquire antibiotic resistance and develop biofilms on both biotic and abiotic surfaces. Recent research has demonstrated that the antidiabetic drug metformin has a potentiation effect on doxycycline and minocycline against certain multidrug-resistant bacterial pathogens, suggesting the potential of this biguanide agent as a novel tetracyclines adjuvant. In this study, we provide evidence showing that the combination of proguanil and chlorhexidine with clarithromycin or rifampicin exhibits rapid bactericidal activities against both planktonic cells and mature biofilms of *A. baumannii*, the capacity to inhibit the development of antibiotic resistance and improvement of the treatment outcomes in *A. baumannii*-infected mice. Given the advantages of repurposing non-antibiotic drugs as antibiotic adjuvants, proguanil and chlorhexidine show promise as adjuvants of specific antibiotics in combating clinically significant pathogenic *A. baumannii*.

## INTRODUCTION

*Acinetobacter baumannii* is a Gram-negative opportunistic pathogen causing hospital- and community-acquired infections affecting various sites, such as the skin, soft tissue, urinary tract, digestive tract, respiratory tract, blood, and central nervous system (1). Despite having lower infection rates compared to other multidrug-resistant (MDR) bacteria (2), *A. baumannii* has become a significant nosocomial pathogen worldwide due to its exceptional genetic plasticity, which enables it to acquire antibiotic resistance traits (3). In the past few years, there has been rapid global dissemination of carbapenem-resistant *A. baumannii* (CRAB) formerly treated with carbapenems as the preferred option for MDR *A. baumannii* infections (4). Currently, polymyxins, tigecycline, and sulbactam are frequently used to treat CRAB infections (5), with polymyxins being the cornerstone of monotherapy and combination therapy despite their systemic toxicities (6, 7). Other antibiotics, *e.g.*, minocycline, trimethoprim-sulfamethoxazole, rifampicin, fosfomycin, macrolide, and aminoglycosides, are rarely used and are typically administered in combination therapy (5). However, colistin (CST) and/or tigecycline-resistant CRAB strains that are refractory to the currently available drugs are increasingly being identified (8). Apart from being pan-drug resistant to last-resort antibiotics, *A. baumannii* poses a challenge in hospital environments due to its ability to form biofilms on both biotic and abiotic surfaces, which contributes to the survival and transmission of *A. baumannii*, and leads to chronic and persistent infections (9). Ventilator-associated pneumonia and catheter-related infections, frequently caused by *A. baumannii* biofilms, are extremely resistant to antibiotic therapy and present a significant clinical management challenge (10). The increasing number of antibiotic-resistant isolates and limited treatment options have prompted researchers to explore alternative approaches to fight against *A. baumannii*-associated infections.

Combination therapy with two or more antimicrobial drugs is a common and critical approach to combat MDR infections (11), which not only shows promise in increasing bactericidal efficacy and slowing down the acquisition of drug resistance but also provides a bet-hedging mechanism to increase the chances of using an effective antibiotic when the pathogen or susceptibility is unknown (12). In addition to combining first-line antibiotics, repurposing non-antibiotic drugs (a.k.a. antibiotic adjuvants) that have undergone extensive toxicological and pharmacological analysis is also an effective approach to reduce the time, cost, and risks of routine antibiotic innovation (13). Combinations of multiple drugs with different modes of action (MOA) and/or toxicity profiles against bacterial planktonic cells and biofilms have been successful in resolving MDR infections (14, 15).

Biguanides are an important class of multifunctional compounds with high therapeutic potential and diverse MOA suitable for the treatments of various pathologies, including antibacterial, anticancer, antidiabetic, antifungal, antimalarial, antiviral, as well as other biological activities, and have regained the spotlight in recent years (16, 17). It has been reported that metformin restores the susceptibility of tetracyclines against MDR *Staphylococcus aureus*, *Enterococcus faecalis*, *Escherichia coli*, and *Salmonella enteritidis*, highlighting the potential of biguanide agents as potential adjuvants for antibiotics (18). Proguanil (PRG), a biguanide derivative, is commonly used in combination with atovaquone as a first-line drug for the prevention and treatment of drug-resistant *Plasmodium falciparum* malaria (19). Chlorhexidine (CHX), another biguanide derivative containing dichloride, is a broad-spectrum cationic antimicrobial agent that disrupts cellular integrity by electrostatically binding to the negatively charged sites of bacterial cell surfaces, which results in bacteriostatic activity at low concentrations and bactericidal activity at higher concentrations (20). Moreover, CHX is also effective at inhibiting and disrupting bacterial biofilms (21). As a result, CHX has become the mainstay topical biocide used clinically for oral disease management, preoperative vaginal cleansing, and prevention of healthcare-associated or skin infections (21–23).

Clarithromycin (CLR), a macrolide antibiotic, effectively halts and inhibits RNA-dependent protein synthesis through reversible binding to the 50S ribosomal subunit of the 70S ribosome in susceptible bacteria (24). Rifampicin (RIF) is a semi-synthetic antibiotic derivative of rifamycin SV, exerting bactericidal effects by binding to DNA-dependent RNA polymerase at the beta subunit, leading to the inhibition of bacterial RNA synthesis (25). Although CLR and RIF are not typically the primary treatment choices for managing *A. baumannii* infection, this study provides evidence that PRG and CHX significantly enhance the activity of both CLR and RIF against *A. baumannii*, demonstrating their potential as adjuvant therapies for related infections.

## RESULTS

### The effectiveness of drug combinations against *A. baumannii* 19606 planktonic cells

Despite the high substructural similarity between PRG and CHX (Fig. 1A), CHX (minimum inhibitory concentration (MIC) =16 mg/L) was significantly more effective than PRG (MIC=1024 mg/L) and less effective than CST (MIC=2 mg/L) against *A. baumannii* ATCC 19606 planktonic cells (Table 1). At 1×MIC, CHX exhibited rapid bactericidal activity, reducing viable cell counts below detection levels within 8 h, similar to CST as a positive control, while PRG alone displayed a slower antibacterial effect (Fig. 1B).

**FIG 1.**
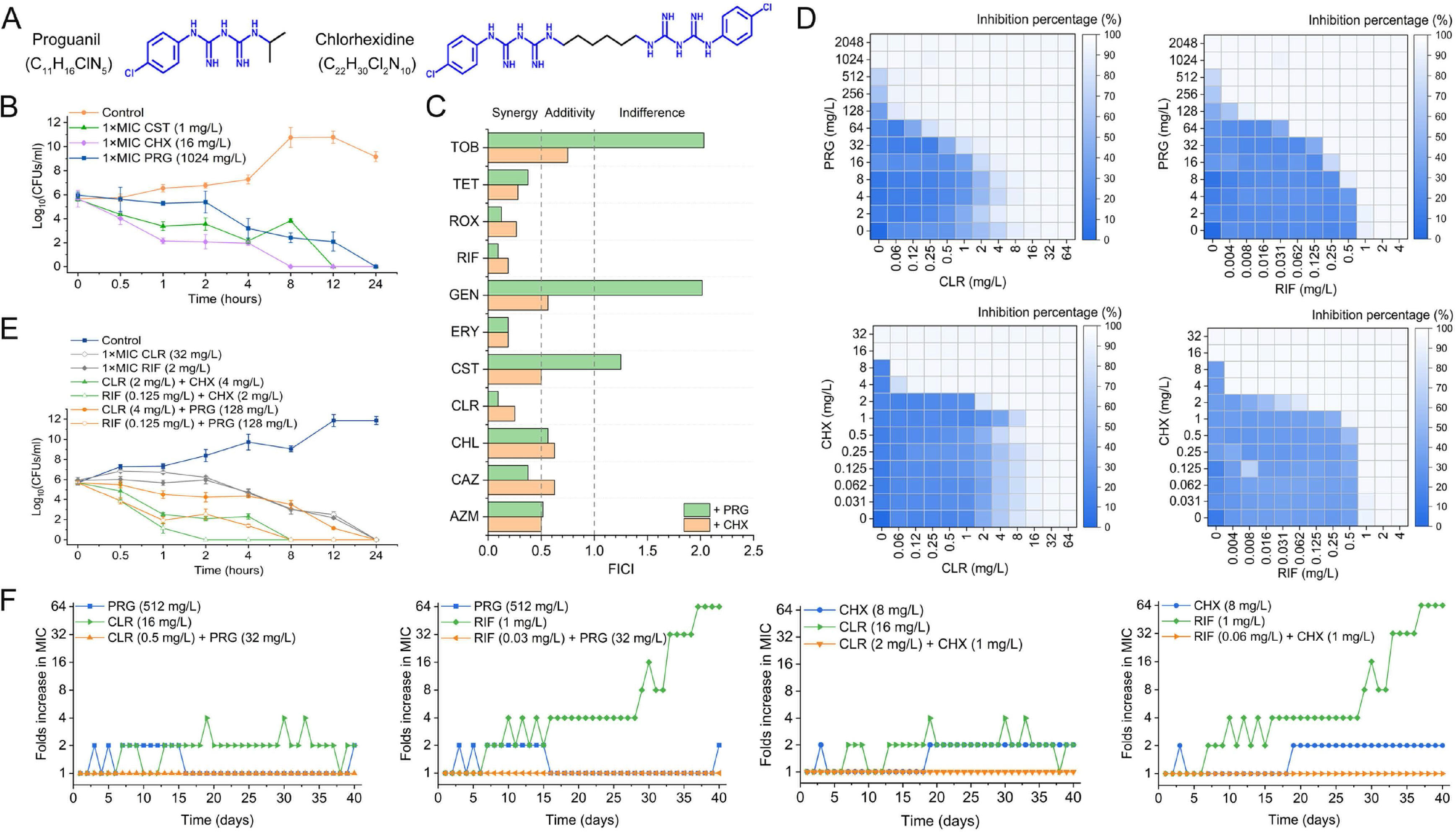
Activities of CLR and RIF in combination with PRG or CHX against *A. baumannii* ATCC 19606 planktonic cells. (A) Chemical structures of PRG and CHX. Blue color highlights similar sub-structures. (B) Time-kill curves of PRG and CHX at 1×MIC. CST is used as a positive control. (C) Activities of different antibiotics in combination with PRG or CHX are assessed using the fractional inhibitory concentration index (FICI). Synergy is defined as FICI ≤ 0.5, additivity as 0.5 < FICI ≤ 1, and indifference as FICI >1. (D) Synergistic growth inhibitory activity of CLR and RIF in combination with PRG or CHX. The color bar represents the corresponding inhibition percentage. (E) Time-kill curves of CLR and RIF alone at 1×MIC or in combination with PRG or CHX. (F) The addition of PRG or CHX prevents the evolution of CLR and RIF resistance to 19606 planktonic cells *in vitro*. The fold increase in MIC after serial passages was determined in the presence of sub-inhibitory concentrations (0.5×MIC) of antibiotics. AZM, azithromycin; CAZ, ceftazidime; CHL, chloramphenicol; CLR, clarithromycin; CST, colistin; ERY, erythromycin; GEN, gentamicin; RIF, rifampicin; ROX, roxithromycin; TET, tetracycline; TOB, tobramycin.

**Table 1.** The MIC (mg/L) and MBC (mg/L) results of single and combination drug treatments against the *A. baumannii* strains.

| *A. baumannii* Strain |  | CST | TGC | MEM | CLR | RIF | PRG | CHX | PRG+CLR | PRG+RIF | CHX+CLR | CHX+RIF |
| --- | --- | --- | --- | --- | --- | --- | --- | --- | --- | --- | --- | --- |
| **ATCC 19606** | MIC | 1 | 0.125 | 0.5 | 32 | 2 | 1024 | 16 | 64 + 1 | 64 + 0.0625 | 2 + 4 | 2 + 0.125 |
|  | MBC | 1 | 0.125 | 0.5 | 32 | 2 | 1024 | 16 | 128 + 4 | 128 + 0.125 | 4 + 2 | 2 + 0.125 |
| **ATCC 17978** | MIC | 1 | 0.125 | 0.0625 | 32 | 2 | 512 | 8 | 32 + 4 | 32 + 0.25 | 1 + 2 | 2 + 0.125 |
|  | MBC | 1 | 0.125 | 0.0625 | 32 | 2 | 512 | 8 | 32 + 4 | 32 + 0.25 | 1 + 2 | 2 + 0.125 |
| **P47** | MIC | 2 | 1 | 16 | 512 | 4 | 512 | 16 | 128 + 32 | 64 + 0.25 | 2 + 64 | 4 + 0.25 |
|  | MBC | 2 | 1 | 16 | 512 | 4 | 512 | 16 | 128 + 128 | 64 + 0.5 | 4 + 16 | 4 + 0.25 |
| **P60** | MIC | 2 | 0.25 | 32 | 256 | 2 | 1024 | 8 | 256 + 8 | 64 + 0.0625 | 2 + 2 | 2 + 0.0625 |
|  | MBC | 2 | 0.25 | 32 | 256 | 2 | 1024 | 8 | 256 + 64 | 64 + 0.125 | 2 + 4 | 2 + 0.0625 |
| **P86** | MIC | 2 | 1 | 64 | 512 | 2 | 512 | 16 | 128 + 64 | 64 + 0.125 | 4 + 8 | 4 + 0.0625 |
|  | MBC | 2 | 1 | 64 | 512 | 2 | 512 | 16 | 128 + 128 | 64 + 0.125 | 8 + 2 | 4 + 0.5 |
| **P95** | MIC | 2 | 0.25 | 64 | 512 | 2 | 512 | 16 | 128 + 32 | 64 + 0.125 | 4 + 8 | 4 + 0.015625 |
|  | MBC | 2 | 0.25 | 64 | 512 | 2 | 512 | 16 | 256 + 128 | 64 + 0.125 | 4 + 32 | 4 + 0.015625 |
| **07AC366** | MIC | >128 | 1 | 64 | 256 | 8 | 1024 | 16 | 256 + 32 | 128 + 0.5 | 4 + 32 | 2 + 1 |
|  | MBC | >128 | 1 | 64 | 256 | 8 | 1024 | 16 | 256 + 32 | 128 + 1 | 4 + 64 | 2 + 1 |
| **978A223** | MIC | 16 | 0.125 | 0.0625 | 256 | 2 | 512 | 8 | 128 + 64 | 128 + 0.0625 | 2 + 64 | 2 + 0.25 |
|  | MBC | 16 | 0.125 | 0.0625 | 256 | 2 | 512 | 8 | 128 + 64 | 128 + 0.125 | 2 + 64 | 2 + 0.25 |
| **978A21** | MIC | 16 | 0.125 | 0.0625 | 32 | 2 | 512 | 8 | 64 + 2 | 64 + 0.25 | 2 + 2 | 2 + 0.5 |
|  | MBC | 16 | 0.125 | 0.0625 | 32 | 2 | 512 | 8 | 64 + 4 | 64 + 0.25 | 2 + 4 | 2 + 1 |

CST, colistin; TGC, tigecycline; MEM, meropenem; CLR, clarithromycin; RIF, rifampin; PRG, proguanil; CHX, chlorhexidine.

To assess synergistic activity, a microdilution checkboard assay was conducted using PRG or CHX combined with 11 commonly used antibiotics with different structural types. As shown in Fig. 1C, both PRG and CHX demonstrated synergistic or additive inhibition of 19606 cell growth with most tested antibiotics, based on the calculated fractional inhibitory concentration index (FICI). The combination of PRG or CHX showed the strongest synergistic effect with CLR (0.09375 with PRG, 0.25 with CHX) and RIF (0.09375 with PRG, 0.1875 with CHX). The presence of a sub-lethal amount of PRG (128 mg/L) or CHX (4 mg/L) significantly increased the susceptibility of 19606 cells to CLR (MIC decreased from 32 to 0.12 mg/L) and RIF (MIC decreased from 2 to ≤0.004 mg/L) (Fig. 1D). Similar conclusions were drawn regarding minimum bactericidal concentration (MBC) (Table 1). Time-kill curve determination (Fig. 1E) showed that CLR or RIF alone at 1×MIC exhibited slow bactericidal processes, whereas CLR+CHX, RIF+CHX, and RIF+PRG displayed a much more rapid killing response to 19606 cells. Co-treatment with 0.125 mg/L RIF and 2 mg/L CHX resulted in no detectable colonies after 2 h.

To further investigate the effect of PRG and CHX on the development of CLR or RIF resistance in *A. baumannii* 19606 cells, serial passaging was performed at sub-lethal antibiotic concentrations with or without PRG or CHX over a 40-day period. As shown in Fig. 1F, the presence of antibiotics alone led to the emergence of resistant cells with up to a 64-fold increase in MIC values, whereas no resistant mutants were observed in the combination group, indicating that combining PRG or CHX effectively prevented the emergence of antibiotic resistance.

### The effectiveness of drug combinations against *A. baumannii* 19606 biofilms

In addition to evaluating the effect of the combined strategy on 19606 planktonic cells, we also assessed its impact on mature biofilms. As shown in Fig. 2A, PRG, CLR, and RIF exhibited effective inhibition of cell release from mature biofilms at high concentrations (2×MIC) and to a lesser extent at low concentrations (0.5×MIC). However, when used in combination, significant inhibition of cell release was observed even at 0.5×MIC concentrations, indicating synergistic effects. Although CHX alone failed to inhibit cell release at 2×MIC, it displayed synergistic effects when combined with CLR or RIF at 0.5×MIC. Subsequently, we investigated the killing effects of these combinations on mature biofilms at 0.5×MIC. Compared to the untreated biofilm, which mainly consisted of viable cells, treatment with PRG or CLR resulted in a slight increase in the proportion of dead cells in each layer of the biofilm, while CHX or RIF treatment did not cause significant changes in cell viability (Fig. 2B). Notably, the combination treatments exhibited a substantial increase in the percentage of dead cells, reaching nearly 50%. Quantitative analysis using COMSTAT also confirmed that the antibiotic combinations led to a significant increase in the percentage of dead cells in the total biomass of the biofilms (Fig. 2C). Furthermore, the combination of antibiotics markedly reduced the maximum thickness (5.51 ± 0.53 ∼ 6.80 ± 1.78 μm) and roughness (0.0137 ± 0.0001 ∼ 0.0447± 0.0009 Ra*) of biofilms compared to the individual antibiotics (8.33 ± 1.15 ∼ 10.0 ± 0.81 μm for maximum thickness and 0.127 ± 0.0156 ∼ 0.408 ± 0.0344 Ra* for roughness).

**FIG 2.**
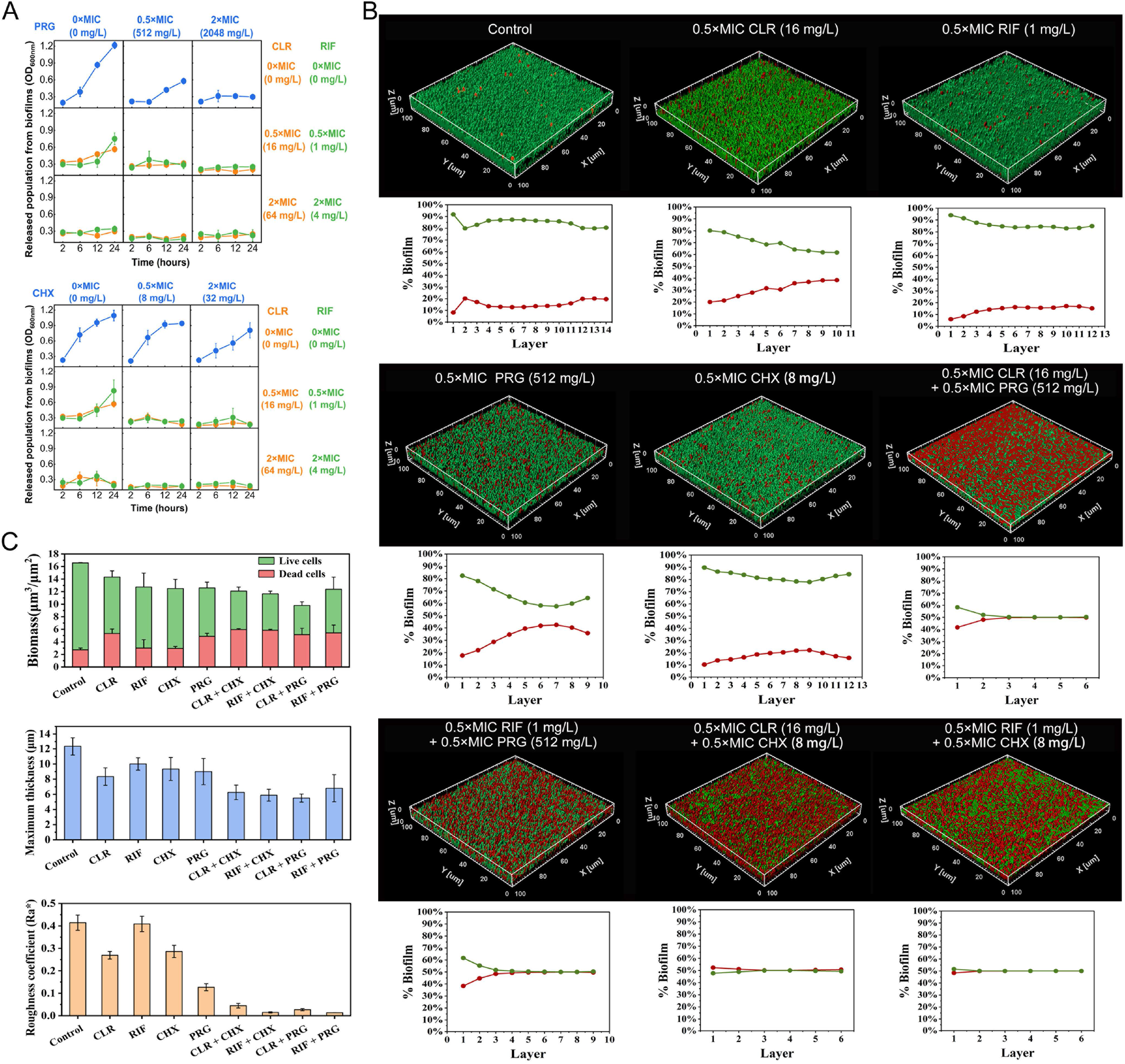
The combined effects of PRG and CHX with CLR or RIF against *A. baumannii* ATCC 19606 mature biofilms. (A) Inhibition of mature biofilm dispersions by PRG and CHX in combination with CLR and RIF at various concentrations: 0×MIC, 0.5×MIC, and 2×MIC. The population released from the biofilms is quantified by measuring the optical density of the supernatant at 600 nm (OD_600 nm_). (B) Bactericidal effects of the PRG and CHX in combination with CLR and RIF at 0.5×MIC on mature biofilms. The samples are compared to untreated samples or those treated with each antibiotic individually. Viable cells are stained with green fluorescence, while dead cells are stained with red fluorescence. The three-dimensional biofilm structures are reconstructed using Imaris software, and the percentages of green and red signals are calculated for each layer. (C) Quantification of biofilm biomass (μm^3^/μm^2^), maximum thickness (μm), and roughness coefficient (Ra*) using ImageJ software with the COMSTAT plugin. Data are reported as mean ± standard deviation.

### The effectiveness of drug combinations against *A. baumannii* clinical isolates

According to the homology to the type strain ATCC 19606 and 17978, seven clinical isolates were identified as *A. baumannii* strains based on their 16S rRNA gene sequence (Fig. S1). Table 1 summarizes the MIC and MBC results of the antibiotics used alone or in combination against these *A. baumannii* strains. All clinical isolates were determined to be MDR based on their susceptibilities to different antibiotics. The MIC values of antibiotics used alone were equivalent to the MBC values. The MIC values ranged from 512 to 1024 mg/L for PRG, 8 to 16 mg/L for CHX, 32 to 512 mg/L for CLR, and 2 to 8 mg/L for RIF. The MIC values of antibiotics used in combinations, calculated based on the lowest FICI, ranged from 32 to 256 mg/L PRG with 1 to 128 mg/L CLR, 32 to 128 mg/L PRG with 0.0625 to 0.5 mg/L RIF, 1 to 4 mg/L CHX with 2 to 64 mg/L CLR, and 2 to 4 mg/L CHX with 0.015625 to 1 mg/L RIF. These values were slightly lower than their respective MBC values.

Subsequently, the combined agents were applied to the mature biofilms of *A. baumannii* clinical strains at 2×MIC. When used individually, the metabolic inhibition rates for the clinical strains ranged from a median value of 27.7% for CLR to 44.0% for PRG (Fig. 3A and S2A). However, when used in combination, the metabolic inhibition rates dramatically increased, ranging from the median value of 82.5% for RIF+CHX to 91.5% for CLR+PRG (Fig. 3B and S2B). The rates of PRG with RIF or CLR were higher (above 90%) than that of CHX with RIF or CLR (above 80%). Furthermore, single-drug administration resulted in lower biomass removal rates for mature biofilms, ranging from a median value of 21.4% for CLR to 30.6% for PRG (Fig. 3C and S2C). Compared to the concentrated metabolic inhibition rates of combination therapy (64.8 to 97.0%), the biomass removal rates of combination therapy showed a distinct variation (8.35 to 89.45%) among all tested strains (Fig. 3D), ranging from more than 80% for strain P60 to less than 50% for strain P47 (Fig. S2D). The efficiency of the four combinations consistently exhibited a high range of 70% to 95% in inhibiting biofilm metabolism, even for the challenging biomass removal strain P47 (Fig. S2B and D).

**FIG 3.**
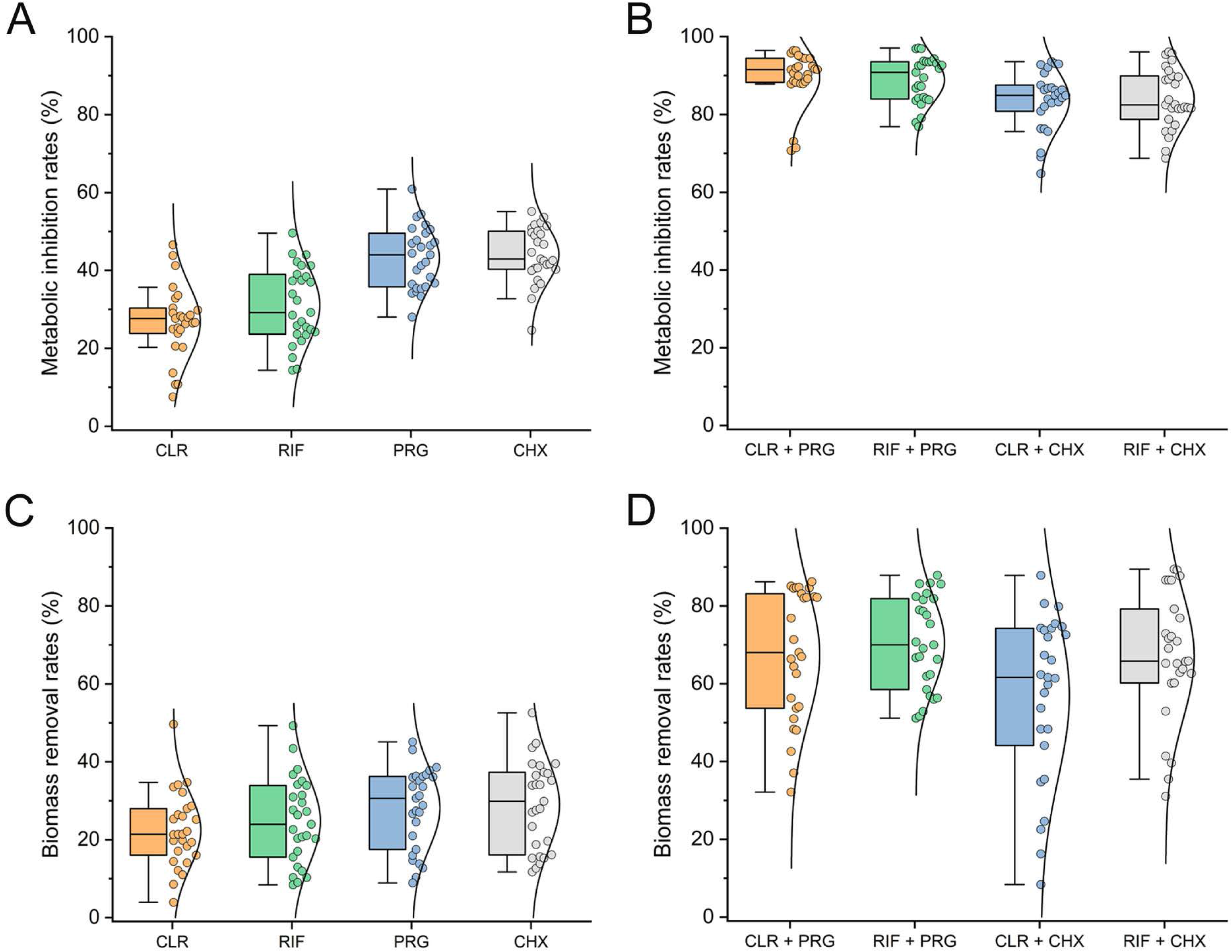
Combined effects of PRG and CHX with CLR or RIF against mature biofilms of *A. baumannii* clinical isolates. Boxplots depicting the variation in metabolic inhibition (A and B) and biomass removal rates (C and D) of CLR, RIF, PRG, and CHX when used alone (A and C) or in combination (B and D) at a final concentration of 2×MIC for the clinical strains. The boxplots show median values and interquartile ranges.

### Drug combinations improve treatment outcomes in *A. baumannii*-infected mice

The efficacy of the combination therapy strategy was evaluated in mice to assess its anti-infective effects. Considering the systemic anti-infective properties of PRG and the topical disinfection capabilities of CHX, two methods of *A. baumannii*-induced intraperitoneal and skin infections in mice were used for the evaluation, respectively. As shown in Fig. 4A, all mice succumbed within 24 h after injection with 1 × 10^8^ CFUs *A. baumannii* ATCC 19606. When 1 × 10^7^ CFUs were injected, the survival rate dropped to 44.4% at 36 h and reached 0% at 48 h. Injection of 1 × 10^6^ CFUs resulted in delayed mortality, occurring at 96 hours. In contrast, all mice in the control group survived until the end of the monitoring period. Based on these results, the injection of 1 × 10^7^ CFUs was deemed an optimal dose with an appropriate latency period and high mortality rate. In comparison to the 50% survival rate observed in the CST group after 96 h, the monotherapy group exhibited a survival rate of only 25% for RIF and 12.5% for PRG and CLR (Fig. 4B). In contrast, the survival rate at 96 h increased to 50% when mice were treated with combination therapy, which was comparable to the positive control CST group.

**FIG 4.**
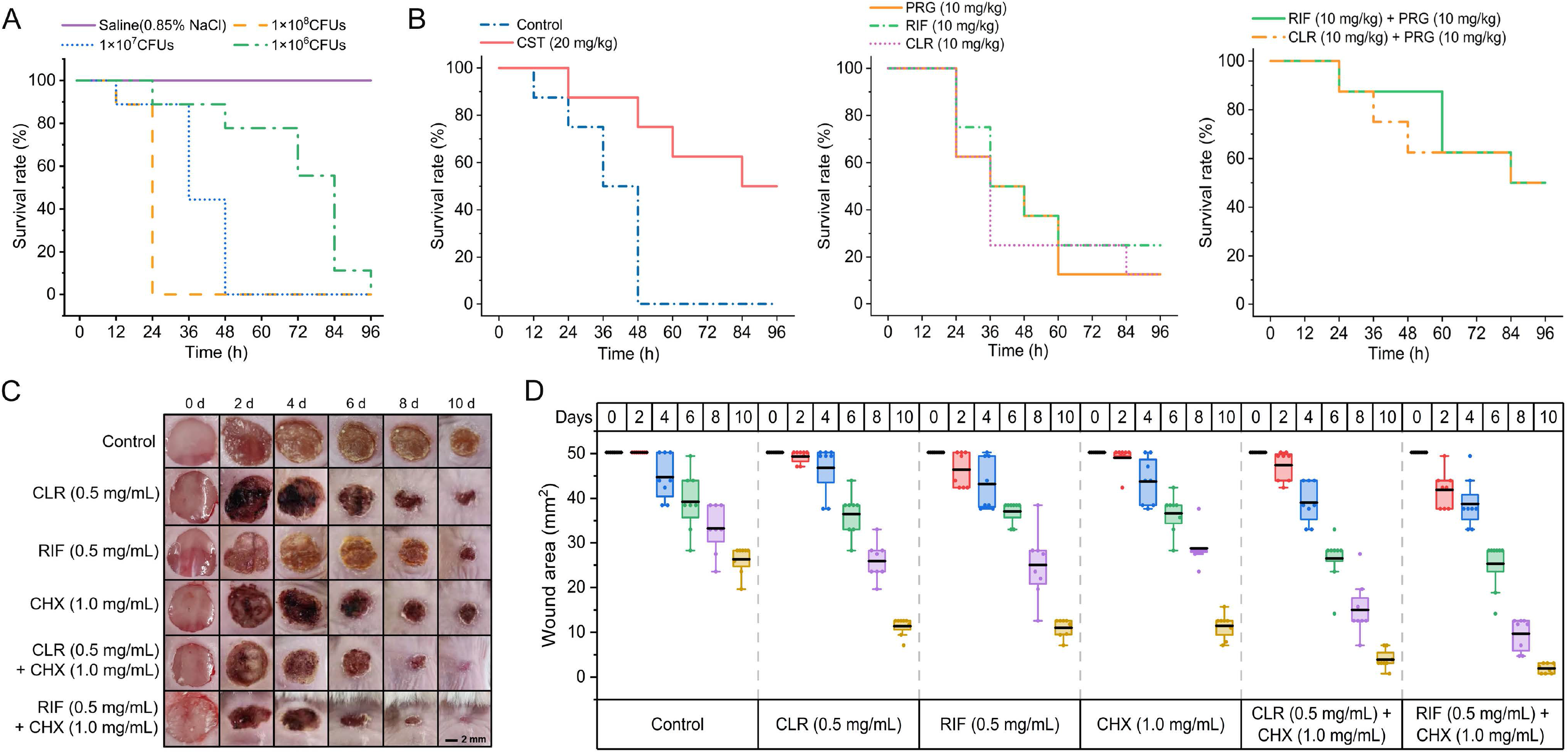
Therapeutic effects of PRG and CHX combined with CLR or RIF on *A. baumannii*-induced intraperitoneal and skin infections in the mice. (A) Survival rates of BALB/C mice infected with various doses of *A. baumannii* ATCC 19606 in the presence of mucin (n = 9). Saline solution was injected as the control group. (B) Survival rates of mice infected with 1 × 10^7^ CFUs 19606 following treatments with PRG (10 mg/kg), RIF (10 mg/kg), CLR (10 mg/kg) alone or in combination (n = 8). CST (20 mg/kg) was used as a positive control. (C) Evaluation of the effects of CHX (1.0 mg/mL), RIF (0.5 mg/mL), and CLR (0.5 mg/mL) alone or in combination on skin wound healing in mice infected with 1 × 10^8^ CFUs/mL 19606. Representative photographs were taken at 0, 2, 4, 6, 8, and 10 d. The group without antibiotic treatment served as the negative control. Scale bar represents 2 mm. (D) Box plots representing the measured wound areas (mm^2^) at 0, 2, 4, 6, 8, and 10 d (n = 8). Each dot represents the value of an individual measurement, and the mean value is shown by the black line.

To investigate the impact of CHX in combination with CLR or RIF on wound healing, the wound area was monitored for 10 d after inducing skin infection in mice with *A. baumannii* ATCC 19606. The group receiving drug intervention exhibited accelerated wound healing compared to the control group (Fig. 4C). Furthermore, the combination therapy group demonstrated a visibly faster healing rate compared to the monotherapy group. After inducing skin infection in mice for 10 d, the wound area in the control group decreased from the initial 50.24 mm^2^ to 25.46 ± 3.47 mm^2^ (Fig. 4D). At this time point, the wound area in the monotherapy group reduced to 11.02 ± 2.24 ∼ 11.68 ± 2.95 mm^2^, while in the combination therapy group, it decreased to 1.96 ± 1.11 ∼ 3.63 ± 2.41 mm^2^. These *in vivo* results demonstrate the potential of PRG and CHX in combination with CLR or RIF as a promising therapeutic strategy for combating bacterial infectious diseases caused by *A. baumannii*.

### PRG and CHX enhance intracellular antibiotic accumulation by increasing bacterial membrane permeability

To further understand the synergistic mechanism of PRG and CHX in combination with CLR and RIF, we investigated their effects on promoting intracellular antibiotic accumulation in *A. baumannii* cells. Due to the poor water solubility of CLR and RIF, as well as the efficient permeation barrier of the gram-negative bacterial outer membranes (OM) to hydrophobic antibiotics (26), we hypothesized that PRG and CHX could enhance the intracellular accumulation of CLR and RIF. To validate this hypothesis, we initially evaluated the effects of PRG and CHX alone or in combination with CLR and RIF on the cell membranes of *A baumannii* ATCC 19606.

Using DiSC_2_(3) as a fluorescent probe, we observed a significant reduction in the red/green fluorescent ratio in cells treated with the combined strategy, indicating a disruption of bacterial membrane potential similar to cells treated with CCCP (positive control) (Fig. 5A). PRG and CHX treatment alone also caused membrane potential dissipation, whereas CLR and RIF treatment did not show such an effect. Additionally, the use of Rosup, a positive contrast agent that induces extra reactive oxygen species (ROS) in cells, resulted in a remarkable increase in fluorescence (Fig. 5B). Interestingly, treatment with PRG alone, but not CHX, either alone or in combination, led to an increase in intracellular ROS levels.

**FIG 5.**
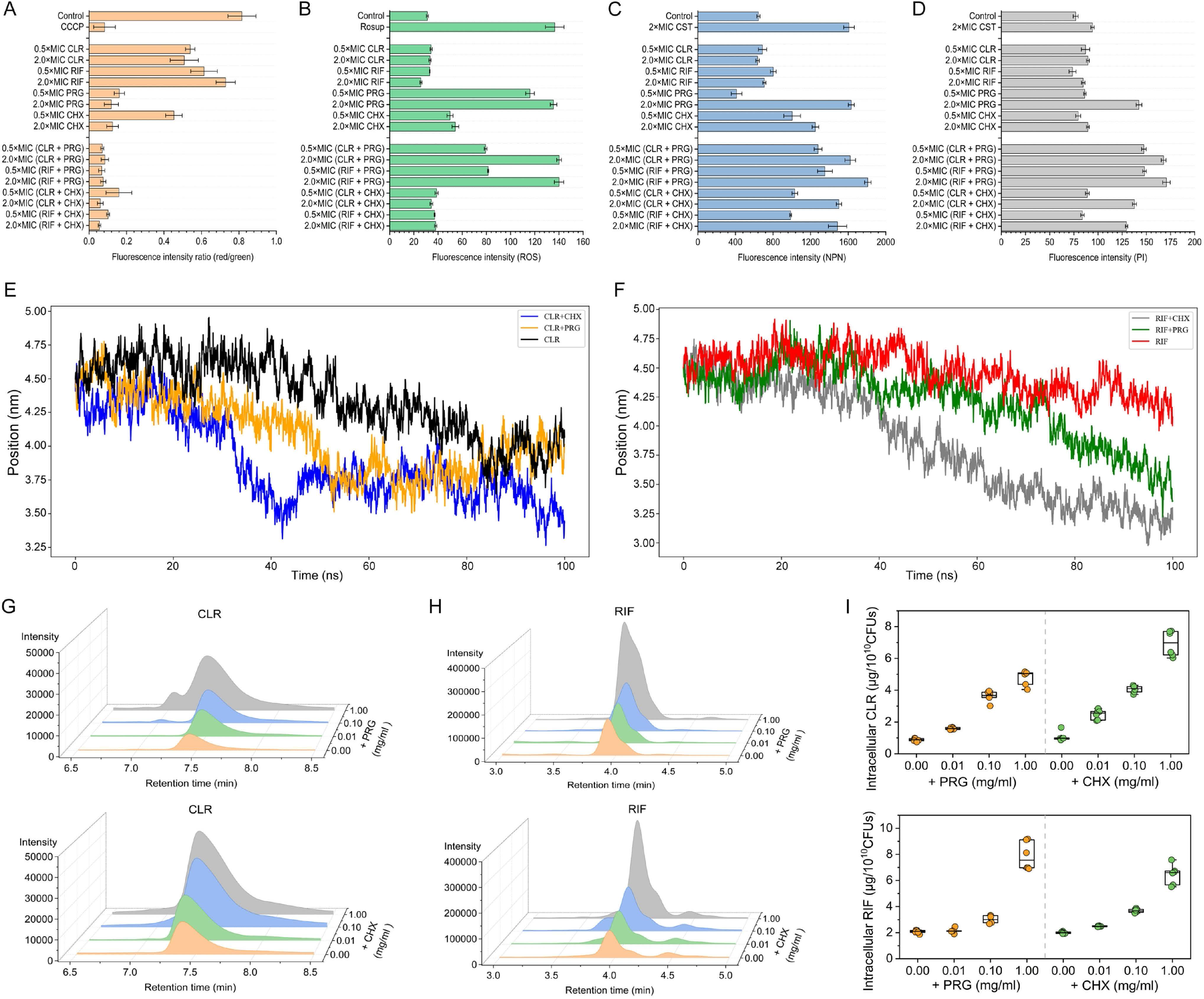
Synergistic mechanisms of PRG and CHX combined with CLR or RIF on *A. baumannii* ATCC 19606. (A) Effects of antibiotics alone or in combination on bacterial membrane potential. CCCP is used as a positive control, showing a significant disruption of bacterial membrane potential based on the reduction in the red/green fluorescence ratio in the DiSC_2_(3)-probed cells. (B) Effects of antibiotics alone or in combination on bacterial ROS production. Treatment with Rosup serves as a positive control for inducing ROS production. (C) Effects of antibiotics alone or in combination on the permeability of bacterial OM by staining with the fluorescent probe NPN. (D) Effects of antibiotics alone or in combination on the permeability of bacterial IM by staining with the fluorescent probe PI. (E) Molecular dynamics simulations of CLR alone or in combination with CHX or PRG crossing the outer membrane in 100 ns. The Y-axis indicates the position of the monitored CLR molecule. (F) Molecular dynamics simulations of RIF alone or in combination crossing the outer membrane in 100 ns. The Y-axis indicates the position of the monitored RIF molecule. (G) HPLC profiles showing the intracellular levels of CLR in bacterial cells treated with PRG and CHX at various concentrations. (H) HPLC profiles showing the intracellular levels of RIF in bacterial cells treated with PRG and CHX at various concentrations. (I) Quantitative results of intracellular CLR and RIF levels represented by box plots with median values and interquartile ranges.

To assess the effects of antibiotics alone or in combination on the permeability of the OM and inner membrane (IM), we employed the fluorescent probes 1-N-phenyl naphthyl amine (NPN) and propidium iodide (PI). Treatment with CST, known for its selective targeting of lipopolysaccharide and disruption of the bacterial OM, resulted in a significant increase in fluorescence intensity for NPN and a weaker increase for PI (Fig. 5C and D). Similar results were observed when CHX was used alone, whereas the combined treatment with CHX showed enhanced fluorescence intensities for both NPN and PI. The simultaneous increase in NPN and PI fluorescence intensity indicated the disruptive effects of PRG, either alone or in combination, on both the outer and inner membranes.

Moreover, molecular dynamics simulations were conducted to simulate the process of CLR and RIF traversing the outer membrane with or without PRG or CHX (Fig. 5E and F). When co-administered with CHX, CLR and RIF demonstrated notable positional changes on the membrane throughout a 100 ns period, decreasing from an initial distance of 4.50 nm to 3.50 nm and 3.25 nm, respectively. In comparison, when CLR and RIF were individually administered, their positions decreased to 4.00 nm and 4.25 nm, respectively. These observations indicate that the presence of CHX enhances the transmembrane efficiency of CLR and RIF. Similar trends were also observed in the combination group with PRG. Subsequently, the levels of intracellular CLR and RIF, when administered individually or in combination, were quantified using HPLC analysis. As anticipated, it revealed a significant increase in intracellular CLR and RIF levels in a concentration-dependent manner when PRG and CHX were co-administered (Fig. 5G-I). Collectively, these findings suggest that PRG and CHX enhance the intracellular accumulation of CLR and RIF by disrupting the membrane potential and increasing membrane permeability.

## DISCUSSION

Currently, the effectiveness of first-line antibiotic therapies, such as carbapenems, tigecycline, and colistin, is limited due to the emergence of MDR and biofilm-forming *A. baumannii* strains (27, 28), which highlights the urgent need for alternative and innovative therapeutic approaches. The combination of tetracyclines with the hypoglycemic drug metformin has shown promising efficacy against various planktonic MDR bacteria, shedding light on the use of biguanides as antibiotic adjuvants (18). In this study, both CHX (MIC = 8 to 16 mg/L) and PRG (MIC = 512 to 1024 mg/L), as biguanide agents, exhibited significantly better antibacterial activity against the tested *A. baumannii* strains compared to the reported MIC of metformin (≥ 10000 mg/L) (18). Calculation of FICI values revealed that both PRG and CHX exhibited synergistic inhibition of planktonic cell growth of *A. baumannii* 19606 when combined with tetracycline or macrolide antibiotics, such as azithromycin, CLR, erythromycin, and roxithromycin. However, they showed additivity or indifference when combined with aminoglycoside antibiotics like gentamicin and tobramycin. Notably, CHX exhibited bactericidal activity at concentrations as low as 1 to 8 mg/L when combined with CLR or RIF, which was significantly lower than the cytotoxic concentrations on human cells (29). For PRG, effective concentrations in combination therapy ranged from 32 to 128 mg/L, which is within the estimated safe range for human administration (30), although its cytotoxic concentration on human cells remains unreported. Subtherapeutic doses of PRG and CHX not only increased the sensitivity of MDR *A. baumannii* to CLR and RIF, but also inhibited the development of antibiotic-resistant phenotypes. Unlike CLR, RIF readily induced antibiotic resistance even at a low concentration of 0.03 mg/L (Fig. S3). The combination of 0.03 mg/L RIF with 32 mg/L PRG or 0.06 mg/L RIF with 1 mg/L CHX significantly suppressed the development of antibiotic resistance (Fig. 1F). This is consistent with consensus on the advantages of combination therapy against resistant bacterial infections (31). In addition to planktonic cells, the synergistic effects of PRG and CHX with CLR or RIF were also observed against the biofilms formed by MDR *A. baumannii*. Co-administration of drugs at 0.5×MIC effectively suppressed the release of the microbial population from mature biofilms and enhanced the bactericidal activity against bacterial cells within the biofilms. However, achieving 100% clearance and killing rate against mature *A. baumannii* biofilms at 2×MIC was challenging, as they are known to be highly resistant and difficult to eliminate entirely (32). Nonetheless, the combination drug strategies demonstrated more pronounced fluctuations in the clearance rate of mature biofilms.

To assess the *in vivo* anti-infective effects of the combination therapy, *A. baumannii*-induced mice intraperitoneal and skin infections were used for PRG and CHX groups, respectively. The equivalent drug doses for mice were calculated based on human data obtained from online drug information sources (https://www.mims.com/) using an established method (33). CHX was used at a concentration of 1 mg/mL, which is commonly found in mouthwash products (34). Although *A. baumannii* ATCC 19606 is not highly virulent, it still effectively induced skin infections in mice. After inducing 10 d of infection, the combination therapy group exhibited significantly smaller wound areas compared to the monotherapy group. To enhance the virulence of strain 19606 and promote sustained intraperitoneal infections in mice, porcine mucin was employed as a virulence-enhancing agent (35). The 96-hour survival rates of infected mice in the combination therapy group were comparable to those treated with CST and significantly higher than the monotherapy group. These findings highlight the potential of PRG and CHX in combination with CLR or RIF as a promising therapeutic strategy against *A. baumannii* infectious diseases *in vivo*. However, further investigations encompassing a broader range of drug concentrations and infections caused by *A. baumannii* clinical isolates with varying levels of virulence are warranted to fully evaluate their clinical potential.

CHX has been reported to exhibit bacteriostatic effects at low concentrations (0.02 to 0.06%) and bactericidal effects at concentrations above 0.1% (21). In this study, we observed the bactericidal effects of CHX against MDR *A. baumannii* at relatively lower concentrations (8 to 16 mg/L, equivalent to 0.0008 to 0.0016%). The antibacterial activity of CHX involves ROS-dependent damage to the bacterial membrane and alterations in membrane compositions (36), which were also observed in *A. baumannii*. Furthermore, CHX was found to increase the permeability of the *A. baumannii* OM, while the permeability of the IM remained unaffected. While PRG and CHX share a similar mechanism of action, PRG was able to concurrently enhance the permeability of both OM and IM of *A. baumannii* cells, which likely contributed to the higher levels of ROS production observed with PRG compared to CHX. Functioning as a prodrug, PRG is converted to the active metabolite cycloguanil by liver cytochrome P450 enzymes and acts as an antifolate agent by inhibiting plasmodial dihydrofolate reductase (37). Currently, there is limited literature discussing the antibacterial mechanisms of PRG and cycloguanil. Although we demonstrated the antibacterial activity of PRG and proposed a possible mechanism for its promotion of antibiotic bactericidal activity *in vitro*, further investigations are required to explore the potential antibacterial activity and synergy of cycloguanil with antibiotics *in vivo*.

In summary, both PRG and CHX enhance the intracellular accumulation of CLR and RIF by increasing the permeability of the bacterial cell membranes, which leads to an increased sensitivity of *A. baumannii* to CLR and RIF. Therefore, the combination of PRG and CHX with CLR or RIF exhibits rapid bactericidal activity against planktonic *A. baumannii*, effectively kills and removes mature biofilms, inhibits the development of antibiotic resistance, and improves treatment outcomes in *A. baumannii*-infected mice. Considering their established safety profiles, PRG and CHX show promise as adjuvants to CLR and RIF for combating clinically significant pathogenic *A. baumannii*. Further prospective clinical trials are necessary to validate the potentiation activity of PRG and CHX with CLR and RIF *in vivo*.

## MATERIALS AND METHODS

### Bacterial strains and antibiotics

All bacterial strains used in this study were listed in Table 1. The nearly full-length 16S ribosomal RNA gene sequences of *A. baumannii* clinical isolates were confirmed by PCR amplification with primer pairs 27F and 1492R (38). *A. baumannii* lab strains ATCC 19606 and ATCC 17978, along with seven clinical strains isolated from Qilu Hospital of Shandong University were routinely grown at 37°C in Tryptone Soy Broth (TSB, Beijing Solarbio Science & Technology Co., Ltd., China). All antibiotics used in this study were obtained from commercial sources.

### Antibacterial activity analysis

Minimum inhibitory concentrations (MICs) of all antibiotics were determined by broth microdilution assay according to the Clinical and Laboratory Standards Institute (CLSI) recommendations for *A. baumannii* (39). Briefly, antibiotics were 2-fold diluted in Mueller-Hinton Broth (MHB, Solarbio, China) mixed with bacterial suspensions containing approximately 5 × 10^5^ colony forming units (CFUs)/mL in a sterilized 96-well microliter plate (NEST Biotechnology, China). After 18 h incubation at 37 °C, the MIC values were defined as the lowest concentrations of drugs with no visible growth of bacteria. After the MIC determination, 50 μL aliquots from all the wells which showed no visible bacterial growth were seeded on TSB agar plates and incubated at 37 °C for 24 h. Minimum bactericidal concentration (MBC) was recorded as the lowest concentration killing 99.9 % of the bacterial population (40). Time-kill curves were performed in triplicate by inoculating 5 × 10^5^ CFUs/mL of the *A. baumannii* 19606 into fresh MHB. Drugs were added individually or in combination at a concentration of 1×MIC. Aliquots were taken at 0, 0.5, 1, 2, 4, 8, 12 and 24 h after inoculation, serially diluted and plated on TSB agar plates to count viable colonies.

### Combined antibacterial activity assay

A microdilution checkerboard method was used to determine the potential effects of individual compound combinations (41). The interactions were evaluated using the fractional inhibitory concentration index (FICI). The FICI was defined as (MIC_A in combination_ /MIC_A alone_) + (MIC_B in combination_ /MIC_B alone_). The interaction inferred from the resulting FICI values was assessed according to the following criteria (42): synergy, ≤0.5; additivity, >0.5 to ≤1; indifference (no interaction), >1 to ≤4; antagonism, >4.

### Antibiotic resistance development assay

To determine the antibiotic resistance development, 5 × 10^5^ CFUs/mL of the *A. baumannii* 19606 at exponential phase were inoculated into fresh MHB medium supplement with antibiotics individually or in combination at a concentration of 0.5 × MIC. After incubation at 37°C for 24 h, a new MIC for bacterial culture was determined as described above. Then, the culture was adjusted to 5 × 10^5^ CFUs/mL and exposed to a new 0.5 × MIC antibiotic environment for the next passage. The process was repeated for 40 d, and the fold increase in MIC of antibiotics relative to the initial MIC was calculated.

### Biofilm formation and quantification

The biofilms of *A. baumannii* were prepared in flat-bottomed microplates following a previously reported method (43) with modifications. Briefly, the overnight cultures of *A. baumannii* were inoculated into TSB medium at a final concentration of approximately 1 × 10^6^ CFUs/mL. Pipet 2.5 mL or 150 μL of adjusted bacterial cultures into each well of 12- or 96-well microplates (NEST Biotechnology, China), respectively. The sterilized glass slides (20 × 20 mm) were inserted vertically into 12-well microplates. Then, 12- or 96-well microplates were incubated at 37 ℃ for 36 h to form biofilms. The mature biofilms formed in 96-well microplates were quantified by staining with crystal violet and measuring optical density (OD) at 590 nm as reported (44). Metabolic activity of living cells in biofilms was assessed by MTT (3-(4,5-dimethylthiazol-2-yl)-2,5-diphenyltetrazolium bromide, Sangon Biotech, China) assay (45).

### Anti-biofilm assay

The non-adherent planktonic cells were removed by gently pipetting the liquid from each well after biofilms formation, and then washed three times carefully with sterile saline (0.85 % NaCl). The antibiotics were diluted in MHB individually or in combination to a final concentration of 0.5 or 2×MIC. In 96-well microplates, 200 μL of each dilution was added to wells containing pre-formed biofilms. The cell proliferation in liquid supernatant was monitored at 37 ℃ for 2, 6, 12, 24 h by measuring OD at 600 nm using a Spectramax 250 microtiter plate reader (Molecular Devices, LLC., USA). To evaluate the removal and killing effects of antibiotics on biofilms, 200 μL of 2×MIC antibiotic dilution was added to pre-formed biofilms and incubated at room temperature for 6 h. After removing the supernatant and gently washing three times, residual biofilms were stained with either crystal violet or MTT as described above for quantitative analysis. The percent reduction (rate) was calculated by the reported equation (46). To visualize biofilm disruption, biofilms grown on glass slides were immersed in 3 mL of antibiotic dilution in each 12-well and incubated at room temperature for 6 hours. Subsequently, the biofilms were stained by LIVE/DEAD^®^ BacLight Bacterial Viability Kit (Invitrogen, Molecular Probes Inc., USA) according to the manufacturer’s instructions and observed by confocal laser scanning microscopy (CLSM). Double-stained cells were viewed under a Zeiss Axio Observer Z1 equipped with a Zeiss LSM700 confocal system. The stack of images in the Z-direction was obtained for three-dimensional images construction. Images representative of the results were reconstructed using IMARIS software (9.0.0, Oxford Instruments, UK). The differences in the structural parameters of the biofilms were confirmed by quantitative analysis using ImageJ software with COMSTAT plugin (47).

### *A. baumannii* infection model

All animal experiments were approved by the Animal Ethics Committee of Shandong University (Permit number: SYDWLL-2023-056). The BALB/c mice ageing 6 to 8 weeks and weighing 16 to 22 g (Jinan Pengyue Experimental Animal. Breeding Co., Ltd., China) were used in the study. Due to the relatively low virulence of *A. baumannii* 19606, porcine mucin was used as a virulence-enhancing agent to promote the development of sustained intraperitoneal infections (35). Cells of 19606 were serially diluted to 2 × 10^6^, 10^7^, and 10^8^ CFUs/mL in sterile saline (0.85 % NaCl). By using a 1-mL syringe, 200 μL-aliquots of each dilution mixed (1:1 v/v) with 10% type II porcine mucin (w/v) (Sigma-Aldrich, China) were injected into the peritoneal cavity of mice (48). The survival trend of mice was monitored for 96 hours.

To establish the skin-infected mice model, the mice were anesthetized with an intraperitoneal injection of 0.7 % sodium pentobarbital (80 mg/kg). After shaving the dorsal hair of mice, depilatory cream was applied to remove any remaining hair (49). The exposed skin was disinfected by applying 75% ethanol solution with a cotton swab, and the residual ethanol was allowed to evaporate naturally at room temperature. Then, a sterile disposable biopsy punch (Rapid Core, Ted Pella, Inc., USA) was used to produce a full-thickness excision wound of 8-mm in diameter on each mouse (50). A sterile cotton swab was soaked in *A. baumannii* ATCC 19606 culture (1 × 10^8^ CFUs/mL) until fully absorbed. The bacterial culture was gently spread onto the skin wound of a mouse using the wet swab and then allowed to air dry at room temperature. All mice were housed in a single cage with sterile paper bedding under well-ventilated conditions in the animal house of Shandong University.

### Antibiotic treatments for infections

Infected mice were randomized into groups (eight mice per group), and antibiotic treatments were initiated 2 h after inoculation. To access the therapeutic efficacy of antibiotics on the intraperitoneal infection model, the agents were administered via intraperitoneal injection with dosages as follows: 20 mg/kg colistin, 10 mg/kg PRG, 10 mg/kg RIF, and 10 mg/kg CLR. An equal volume of sterile saline (0.85 % NaCl) was injected as a negative control group. The survival rates of mice were recorded every 12 hours. For the skin infection model, the wound cavity was gently swabbed every day by a cotton swab fully saturated with antibiotic solutions containing 1 mg/mL CHX, 0.5 mg/mL RIF, or 0.5 mg/mL CLR. The diameters of wound areas were measured in two directions every two days.

### Bacterial membrane potential assay

*A. baumannii* 19606 was diluted to OD_600nm_ of 0.5 by MHB and incubated with antibiotics (0.5 or 2 × MIC) at 37℃ for 1.5 h. Then, the cells were washed and resuspended with phosphate-buffered saline (PBS, pH 7.4). The BacLight™ Bacterial Membrane Potential Kit (Invitrogen, China) was used to measure the bacterial membrane potential according to the manufacturer’s instructions. Stained bacterial cells were assayed in an ImageStream^X^ Mark II imaging flow cytometer (Millipore, USA) to collect the red and green fluorescence intensity.

### Reactive oxygen species assay

Accumulation of intracellular ROS was quantified with a Reactive Oxygen Species Assay Kit (Beyotime, China). Briefly, 0.5 OD_600nm_ of collected *A. baumannii* 19606 cells were resuspended with MHB containing 10 μM DCFH-DA (2′,7′-dichlorodihydrofluorescein diacetate). Subsequently, cells were treated with 0.5 or 2 × MIC antibiotics at 37 °C for 1.5 h in the dark. DCFH is non-fluorescent unless oxidized by intracellular ROS to transform DCF. DCF fluorescence intensity was measured by a SpectraMax Gemini^®^ XPS spectrofluorometer (Molecular Devices, USA) with the excitation source at 488 nm and emission at 525 nm.

### Membrane permeability assay

The 1-N-phenylnaphthylamine (NPN) and propidium iodide (PI) accumulation assay were adapted to measure the permeability of bacterial outer and inner membranes (18). Briefly, after incubation with antibiotics (0.5 or 2×MIC), the *A. baumannii* 19606 cells were washed and resuspended with 5 mM HEPES (pH 7.0) buffer in the presence of 10 μM NPN or PI. After incubation at room temperature for 10 min in the dark, fluorescence intensity was measured using excitation and emission wavelengths of 350 nm and 420 nm for NPN or excitation and emission wavelengths of 490 nm and 635 nm for PI.

### Molecular dynamics simulation

The system consisted of a double-layered membrane with a composition of POPE: POPG: POCL1 = 75%: 25%: 5% and drug molecules (51). A planar region with a thickness of 4.1 nm along the Z-axis was designed within a 10 nm cubic box, followed by the addition of DPPC. The Martini coarse-grained force field (3.0) was used to represent the system components, and the membrane and drug molecules were built using the Charmm-gui Martini Maker web server (52). Molecular dynamics simulations were performed using GROMACS v5.12 in the NPT ensemble at a temperature of 310 K and a pressure of 1 bar for a total of 100 ns (53). Temperature and pressure were coupled using the velocity-rescale method (time constant of 1 ps) and semi-isotropic pressure coupling with the Parrinello-Rahman algorithm (time constant of 5 ps), respectively (54), using the Martini force field. The system was energy-minimized using the steepest descent algorithm. Umbrella sampling was conducted along the bilayer normal (Z-axis) within a range of 0.0 to 4.1 nm, with a step size of 0.1 nm, generating 10 windows. The solute was centered using a harmonic biasing potential (k = 300 kcal/mol/nm^2^). The simulation trajectories were visualized using the Visual Molecular Dynamics (VMD) software for analysis of the structural properties of the membrane and drug molecules (55).

### Antibiotics accumulation detection

An overnight culture of *A. baumannii* 19606 was adjusted to 2 × 10^8^ CFUs/mL by fresh sterile saline. The culture was aliquoted into 50 mL and transferred to 100 mL centrifuge tubes. After centrifugation at 6000 rpm for 5 min, the precipitate was resuspended with a 50 mL solution of either CLR (1 mg/mL) or RIF (1 mg/mL) together with varying concentrations (0, 0.01, 0.1, or 1 mg/mL) of PRG or CHX. After incubation at 37 °C for 1 h, the bacterial cells were pelleted by centrifuging at 4000 rpm for 10 min and then washed twice with sterile saline. The antibiotics accumulated in the cells were extracted with a mixture of acetonitrile and dichloromethane (6:4, v/v). After centrifugation for 10 min at 8000 rpm, the supernatant was transferred to a round flask. This step was repeated three times to thoroughly extract the antibiotics. The obtained extracts were combined and then evaporated to dryness in a rotary evaporator at 40 °C under vacuum. After that, 2 mL acetonitrile: water containing 0.067 M potassium dihydrogen phosphate and 0.014 M triethylamine (pH 5.5) (6:4, v/v) and acetonitrile: methanol: 0.04 M potassium dihydrogen phosphate (35:35:30, v/v/v) were used to dissolve the extracted CLR and RIF, respectively. The obtained samples were analyzed by a Shimadzu LC 20AT HPLC system (Shimadzu, Japan) after filtering through a 0.22 µm membrane. The liquid chromatography separation was performed on a Hypersil™ BDS C18 column (250 × 4.6 mm, 5 μm) with mobile phase acetonitrile: water containing 0.067 M potassium dihydrogen phosphate and 0.014 M triethylamine (pH 5.5) (6:4, v/v) for CLR or acetonitrile: methanol: 0.04 M potassium dihydrogen phosphate (35:35:30, v/v/v) RIF. The flow rate was 1 mL/min. The injection volume was 10 μL. The calibration curves of the standards were used to determine the concentrations of the antibiotics in the extracts. Peak areas were used for quantitative analysis.

## ETHICS STATEMENT

*A. baumannii* isolates were collected and purified from routine diagnoses of Qilu Hospital. No patient-related data were analyzed. According to the regulations of Qilu Hospital of Shandong University, such research does not require ethical approval.

## ACKNOWLEDGMENTS

This work was supported by the Taishan Industry Leading Talent Program (Tscy20200334, to WH), National Natural Science Foundation of China (32070100, to WH), National Key R&D Program of China (2021YFC2101000, to WH), Shandong Provincial Natural Science Foundation (ZR2021QC087, to CW).

The authors would like to thank Zhifeng Li, Haiyan Yu, Yuyu Guo, and Xiangmei Ren of the Core Facilities for Life and Environmental Sciences, State Key Laboratory of Microbial Technology of Shandong University for the technical assistance in operating flow cytometer, CLSM, and HPLC.

---

**FIG S1.**
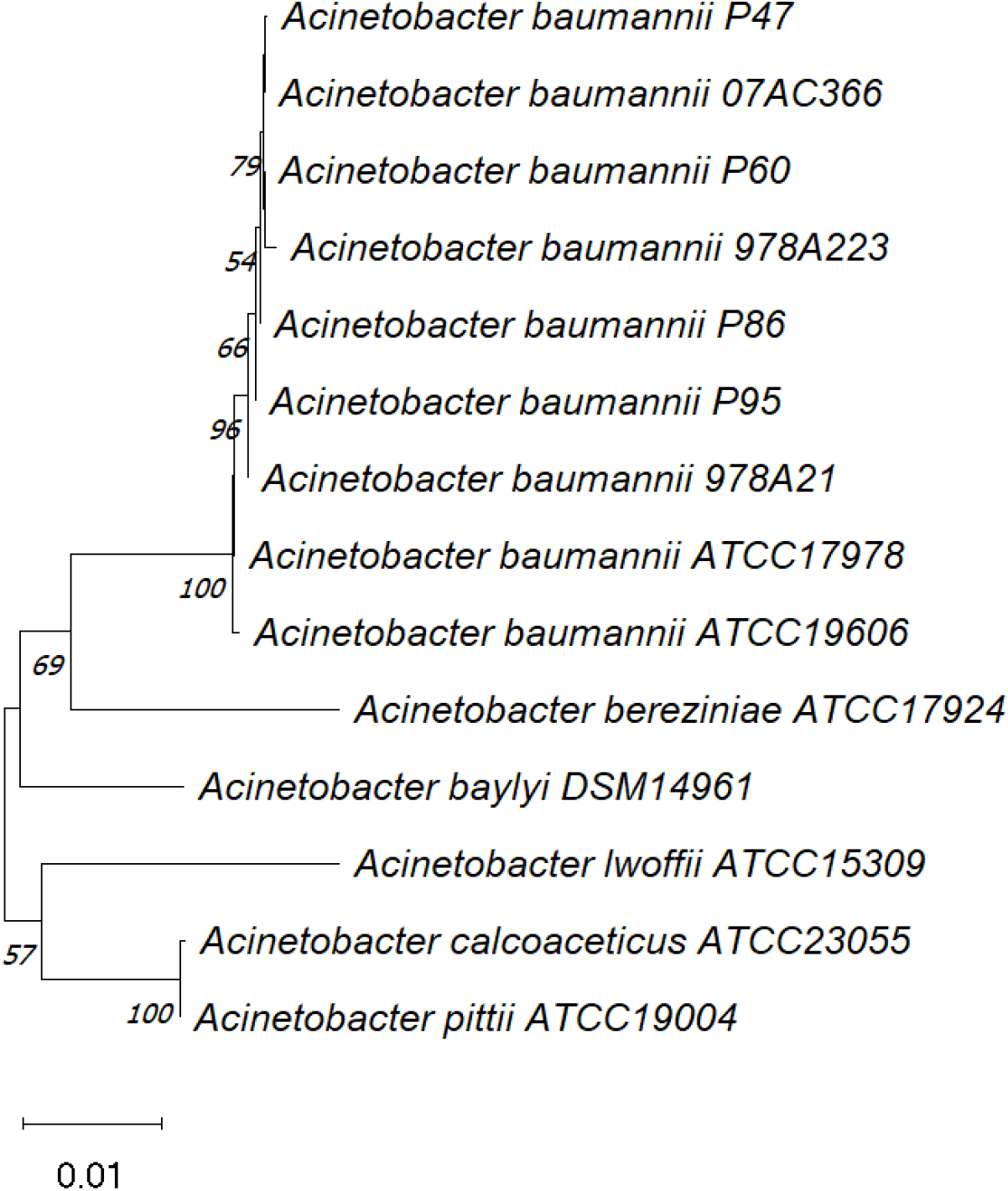
Phylogenetic tree of seven *A. baumannii* clinical isolates based on 16S rRNA gene sequences with homologous sequences from type strain ATCC 19606 and 17978. The 16S rRNA gene sequences of *Acinetobacter baylyi*, *Acinetobacter bereziniae*, *Acinetobacter calcoaceticus*, *Acinetobacter lwoffii*, *Acinetobacter pittii* were used as outgroup taxa.

**FIG S2.**
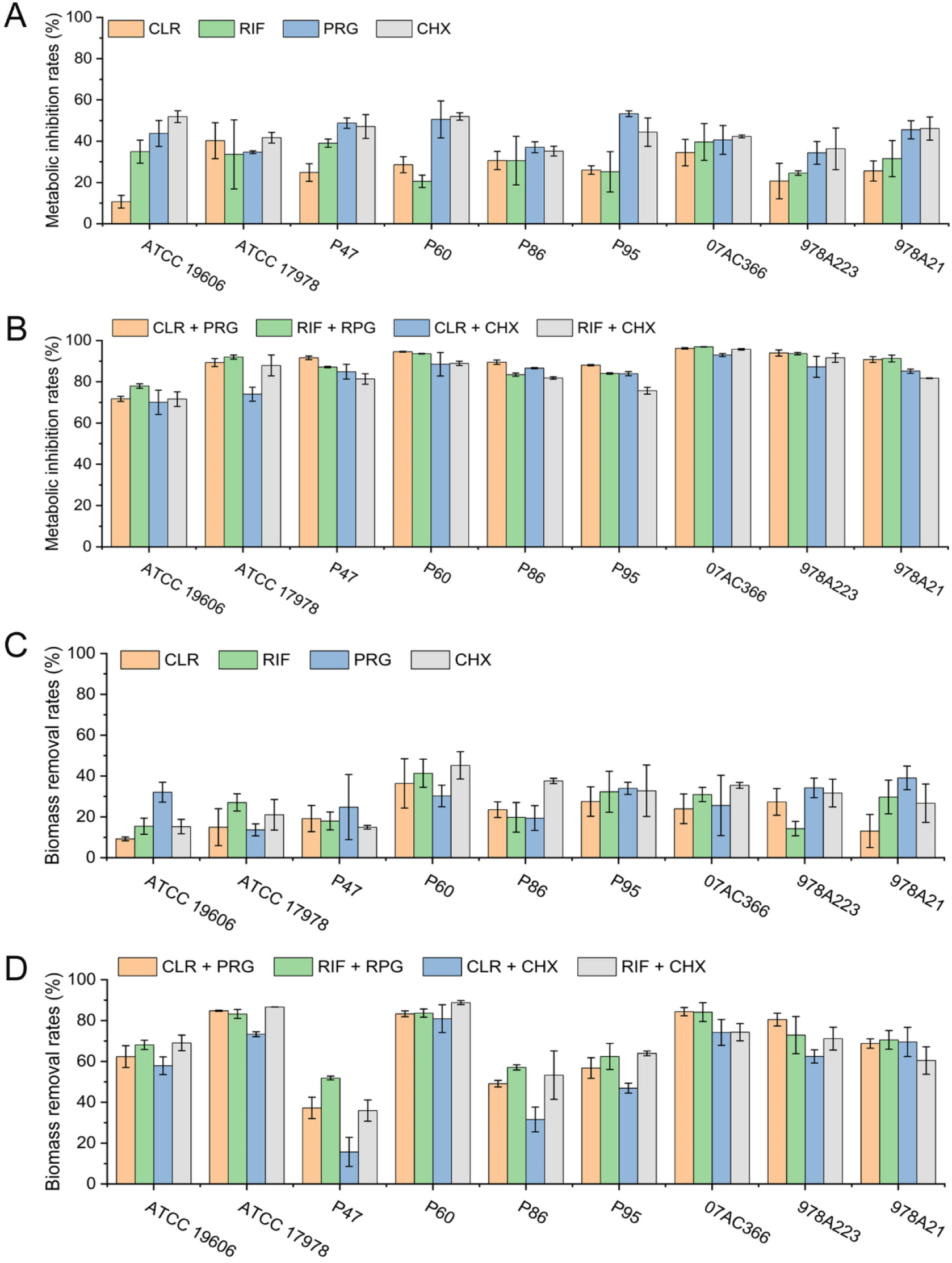
Metabolic inhibition (A and B) and biomass removal rates (C and D) of CLR, RIF, PRG, and CHX when used alone (A and C) or in combination (B and D) at a final concentration of 2×MIC for nine *A. baumannii* strains.

**FIG S3.**
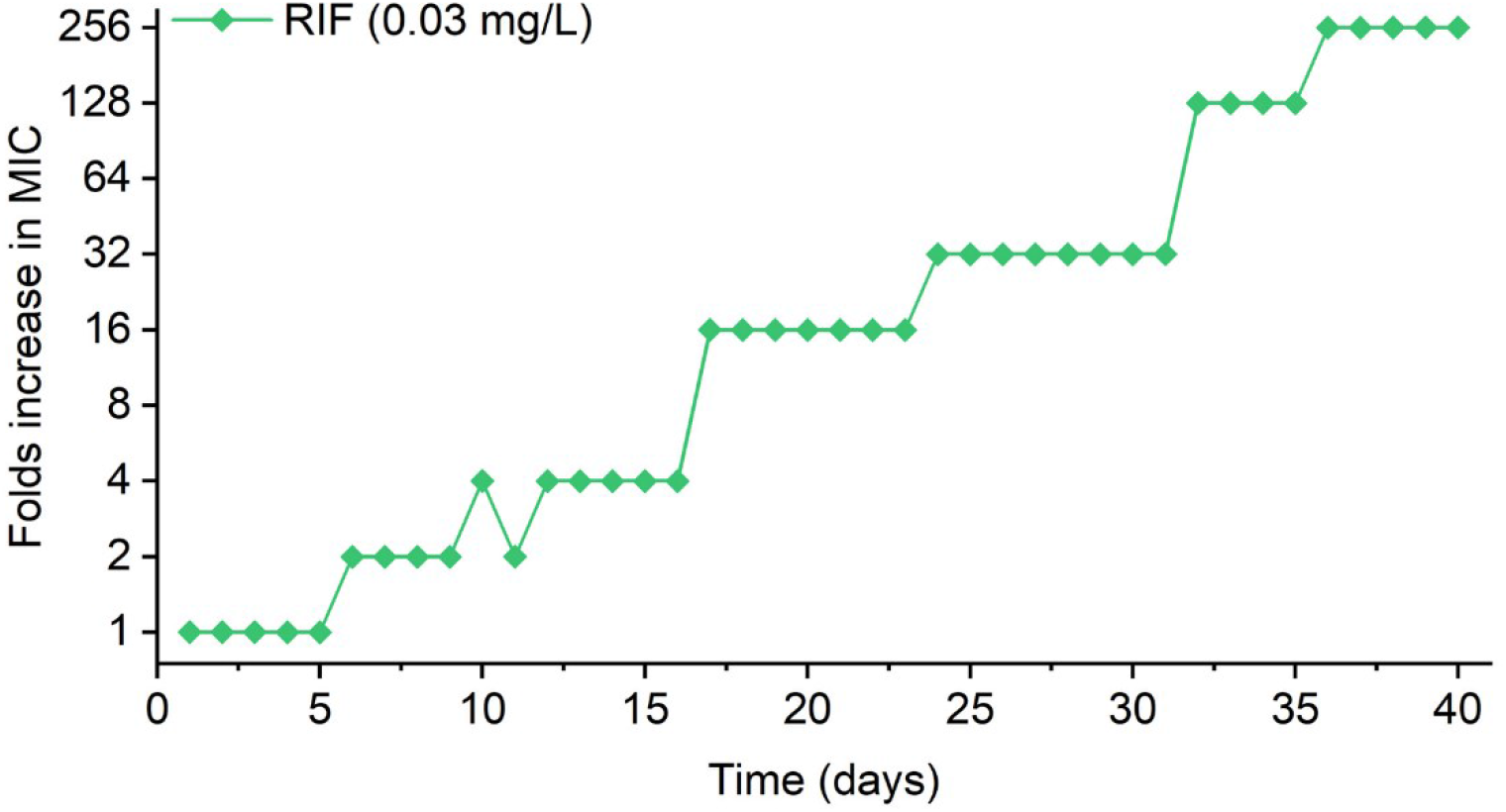
Increase in MIC of *A. baumannii* ATCC19606 after serial passages in the presence of sub-inhibitory concentrations of RIF. The initial concentration of RIF was set as 0.03 mg/L.

